# Synthetic lethal targeting of *TET2*-mutant hematopoietic stem and progenitor cells by XPO1 inhibitors

**DOI:** 10.1101/2022.10.12.511957

**Authors:** Chang-Bin Jing, Nicole Prutsch, Shuning He, Mark W. Zimmerman, Yosef Landesman, A. Thomas Look

**Author notes:** These authors contributed equally to this manuscript (Chang-Bin Jing, Nicole Prutsch). **Corresponding author**: A. Thomas Look, MD, Department of Pediatric Oncology, Dana-Farber Cancer Institute, 450 Brookline Ave, Boston, MA 02115.

## Abstract

*TET2* inactivating mutations serve as initiating genetic lesions in the transformation of hematopoietic stem and progenitor cells (HSPCs). In this study, we analyzed known drugs in zebrafish embryos for their abilities to selectively kill *tet2*-mutant HSPCs *in vivo*, and we found that the exportin 1 (XPO1) inhibitors, selinexor and eltanexor, selectively kill *tet2*-mutant HSPCs. In serial replating colony assays, these small molecules were selectively active in killing murine *Tet2*-deficient Lineage-, Sca1+, Kit+ (LSK) cells, and also *TET2*-inactivated human acute myeloid leukemia (AML) cells. Selective killing of *TET2*-mutant HSPCs and human AML cells by these inhibitors was due to increased levels of apoptosis, without evidence of DNA damage based on increased γH2AX expression. The finding that *TET2* loss renders HSPCs and AML cells selectively susceptible to cell death induced by XPO1 inhibitors provides preclinical evidence of selective activity of these drugs, justifying further clinical studies of these small molecules for the treatment of *TET2*-mutant hematopoietic malignancies and to suppress clonal expansion in age-related *TET2*-mutant clonal hematopoiesis.

## Introduction

TET2 is a member of the TET protein family of proteins, and a critical regulator of DNA demethylation by catalyzing the conversion of 5-methylcytosine (5-mc) to 5-hydroxymethylcytosine (5-hmc) [1]. Through this enzymatic activity, TET2 is thought to initiate the reversal of aberrant cytosine methylation in enhancers and CpG islands, thus maintaining tissue-specific patterns of gene expression [2–4]. TET2 may also play other roles in the control of gene expression and alternative splicing [5]. Interestingly, *TET2* is one of the most frequently mutated tumor suppressor genes in myeloid malignancies and has been identified in 10-25% of patients with acute myeloid leukemia (AML), 10-13% with myeloproliferative neoplasms (MPN), and 20-30% with myelodysplastic syndrome (MDS) [6–11]. This has led to the hypothesis that *TET2* mutations represent initiating mutations in the transformation of hematopoietic stem and progenitor cells (HSPCs), inducing a premalignant state of clonal dominance that predisposes to the acquisition of additional mutations, which then lead to overt hematologic malignancies [12].

*TET2*-inactivating mutations have also been identified in persons over 40 years of age with age-related clonal hematopoiesis (ARCH), also referred to as clonal hematopoiesis of indeterminant potential (CHIP), a “pre-malignant” condition involving aberrant clonal expansion of mutant HSPCs in the bone marrow [13–15]. Patients with ARCH are predisposed to develop hematologic malignancies and the mutant HSPCs also give rise to activated macrophages that invade arterial endothelium and promote atherosclerosis, thus causing an increased risk of heart attacks and strokes [13–15]. Due to the increased risk of mutations arising over time, ARCH becomes more frequent in the elderly population, affecting 5.6% of people over 60 years of age and up to 18.4% of people over 90 years of age [13].

Published studies indicate that mutations of epigenetic tumor suppressors, such as *DNMT3A* and *TET2* occur as transformation initiating events in HSPCs, causing clonal expansion that favors the acquisition of additional mutations and other genomic abnormalities leading to AML, MPN and MDS [16–19]. AML patients harboring these mutations often respond to treatment and patients enter remission. However, the clone harboring the initial inactivating mutation in *DNMT3A* or *TET2* is often still present during morphologic remission in AML patients, providing a clonal advantage for the mutant HSPCs. Studies have shown that these mutant HSPCs then frequently acquire new cooperating mutations in genes such as FLT3-ITD or NPM1c, which cause clonal outgrowth and relapse as AML [20, 21].

Thus, for therapy of myeloid malignancies with *TET2* mutations, there is a need for drugs targeting molecules that are synthetic lethal with *TET2* loss. Such drugs might be especially useful during remission, when it may be possible to eradicate or suppress the replication of persistent mutant HSPCs and prevent their clonal evolution to AML. Drugs that selectively kill *TET2*-mutant HSPCs would also be candidates for *in vivo* studies in murine models to test the hypothesis that administration of selectively active drugs might reverse atherosclerosis and reduce the frequency of heart attacks and strokes in patients with ARCH due to *TET2* mutations.

To identify potential therapies for patients with *TET2*-mutant myeloid malignancy that could be rapidly translated to the clinic, we screened a *tet2*-loss-of-function zebrafish line [22] with a panel of 129 FDA-approved anticancer drugs (AODVII from NCI/NIH). As we reported previously, this study showed that TOP1-targeted drugs and PARP1 inhibitors selectively kill *tet2*-mutant compared to wild-type HSPCs [23]. We showed that this selectivity is because loss of *tet2* led to reduced expression levels of Tyrosyl-DNA phosphodiesterase 1 (*tdp1*), which in turn impaired the ability of *tet2*-mutant cells to repair double-strand breaks induced by TOP1-targeted drugs and PARP1 inhibitors [23]. We went on to show that TDP1 expression levels were reduced in *Tet2* mutant compared to wild-type murine HSPCs and the cells were also more sensitive to TOP1-targeted drugs and PARP inhibitors in serial replating colony-forming assays. Apparently, in the absence of TET2 function, methylation accumulates in the *TDP1* CpG island or other regulatory sequences, leading to the aberrant reduction of expression levels of mRNAs encoding TDP1. However, cells treated with TOP1-targeted drugs and PARP inhibitors undergo DNA damage, which could lead to genomic instability with progression to new myeloid malignancies or secondary myelodysplastic syndrome in patients treated with these drugs.

In our current study, we extended our screen to evaluate the selectivity of drugs not included in our initial libraries, including newer drugs currently in phase I/II clinical trials. We found that among the drugs we evaluated, the XPO1 inhibitors, selinexor and eltanexor, were among the most selective drugs we evaluated in the zebrafish model for killing *tet2*-mutant compared to wild-type HSPCs. These two XPO1 inhibitors have shown activity in Phase I and II clinical trials for patients with AML, and selinexor has been FDA-approved for the treatment of relapsed/refractory multiple myeloma and diffuse large B-cell lymphoma [24–26]. These two drugs incorporate the same warhead for covalent binding to the XPO1 cargo binding pocket and are thought to have similar mechanisms of action. Eltanexor is a second-generation inhibitor with a much lower ability to cross the blood-brain barrier and thus can be given by daily oral dosing with less CNS-mediated nausea and anorexia than selinexor.

Here we show that XPO1 inhibitors are selectively active against *tet2*-mutant HSPCs by examining the number of definitive HSPCs in the caudal hematopoietic tissue of zebrafish embryos with *tet2*-mutations compared to wild-type embryos. We also evaluated murine *Tet2*-deficient HSPCs compared to wild-type HSPCs, by treating lineage-negative, Sca1-positive, and c-Kit-positive (LSK) cells that were grown as colonies in repetitive replating assays in vitro in methylcellulose medium. We found that each of the XPO1 inhibitors specifically reduced colony formation of *Tet2*-mutant LSK cells at concentrations that had little effect on wild-type LSK cells. Importantly, γ-H2AX expression assays did not reveal any evidence of DNA damage in zebrafish HSPCs or human AML cell lines treated with the XPO1 inhibitors. In addition, both selinexor and eltanexor showed selective killing of human AML cell lines after they had been rendered *TET2*-deficient using CRISPR-cas9, supporting the selective activity of XPO1-inhibitors in killing *TET2*-deficient HSPCs.

## Results

### Selinexor and eltanexor selectively kill *tet2*-mutant hematopoietic stem and progenitor cells (HSPCs) at dosages showing little effect on wild-type HSPCs

We studied these two drugs in zebrafish embryos as part of an analysis of approved drugs for selective activity in killing *tet2*-mutant compared to wild-type HSPCs. Thus, *tet2+/+, tet2*+/− and *tet2*−/− embryos were exposed to multiple concentrations of selinexor and eltanexor, by adding one of these drugs or DMSO to the embryo water in 24-well plates starting at 1-day postfertilization (dpf). At 5 dpf, the embryos were fixed with 4% paraformaldehyde (PFA), and effects on HSPC numbers residing within the caudal hematopoietic region (CHT) of the embryo were evaluated by whole-mount in situ hybridization (WISH) with the *c-myb* riboprobe. As shown in **Fig. 1**, after testing multiple drug combinations we found that selinexor (100 nM) and eltanexor (250 nM) selectively killed *tet2*-mutant HSPCs without measurable effects on wild-type HSPCs. Thus, at these concentrations, each of the XPO1 inhibitors is synthetic lethal *in vivo* with *tet2* loss in *tet2*-mutant HSPCs.

**Figure 1.**
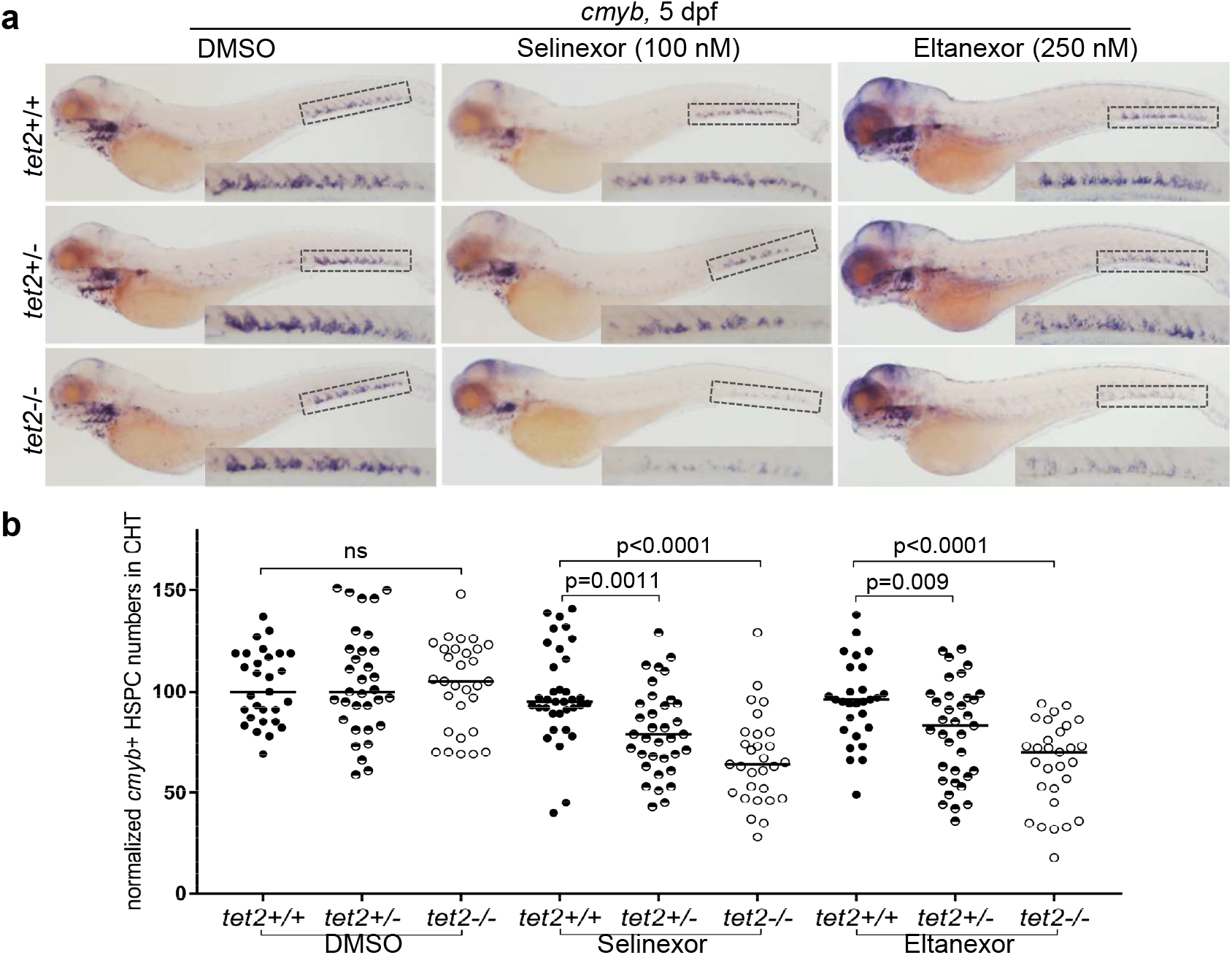
*tet2*-mutant HSPCs are hypersensitive to selinexor and eltanexor. **(a)** Embryos with the indicated genotypes were treated with either DMSO, selinexor, or eltanexor from 1 to 5 dpf. WISH was performed using a riboprobe for *c-myb* at 5 dpf (Overall magnification: 5X). Insets in each of the images show a magnified image of the CHT region (Magnification in the inset: 11.25X). CHT: caudal hematopoietic tissue. (**b**) Quantification of cells exhibiting *c-myb* expression in the CHT region shown in panel a.

### Selinexor and eltanexor treatment do not affect the birth of HSCs from the AGM region

To determine whether the XPO1 inhibitors inhibit the production of HSCs from the ventral wall of the aorta, the zebrafish aorta-gonad-mesonephros (AGM) region, we analyzed the effects of these drugs on the numbers of newly formed HSPCs in the developing zebrafish embryo. In this experiment, HSPCs recently formed from endothelial cells of the ventral wall of the aorta were identified by WISH using the *c-myb* riboprobe at 2 dpf. We found that XPO1 inhibitors administered from 1 to 2 dpf did not affect the generation of HSPCs in the AGM region in either wild-type or *tet2*-mutant HPSCs (**Fig. 2a and b**).

**Figure 2.**
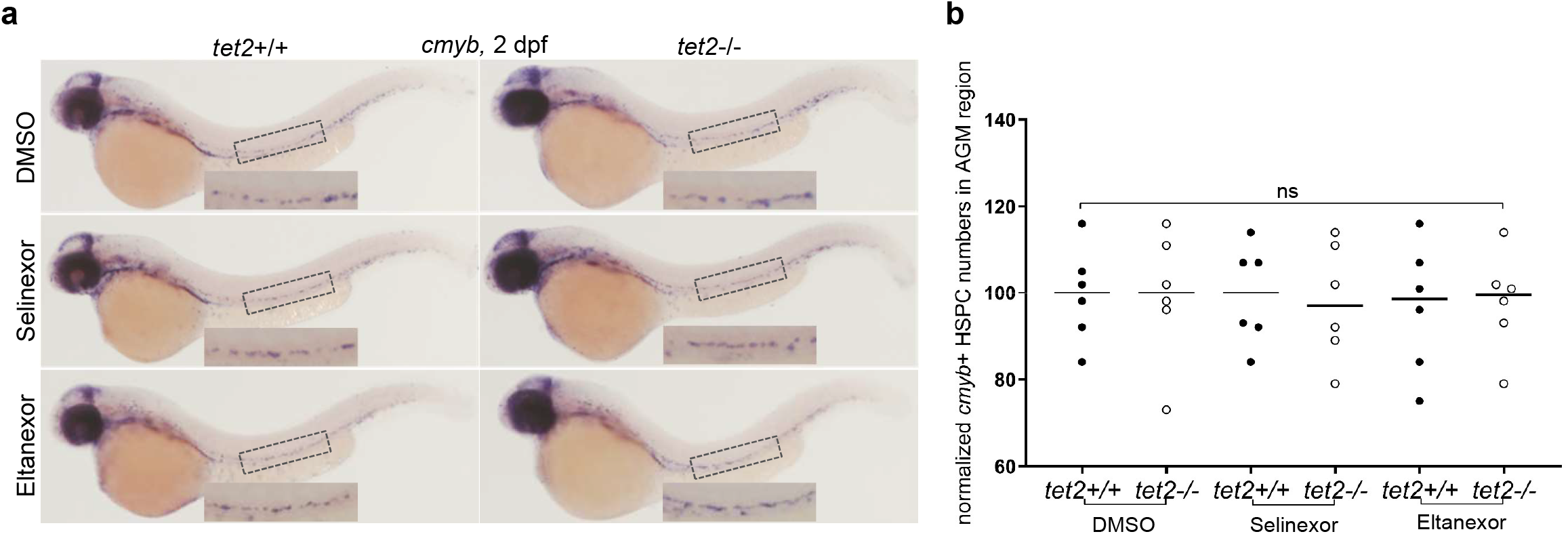
Selinexor and eltanexor treatment don’t affect the birth of HSCs from the AGM region. **(a)** Embryos with the indicated genotypes were treated with either DMSO, selinexor (100 nM), or eltanexor (250 nM) dissolved in fish water from 1 to 2 dpf. WISH was performed using a riboprobe for *c-myb* at 2 dpf and images were taken (Overall magnification: 5X). Insets in each of the image magnify staining in the AGM region (Magnification in the inset: 11.25X). AGM: Aorta-gonad-mesonephros. (**b**) Quantification of cells with *c-myb* expression in the AGM region, as shown in panel a.

### Selinexor and eltanexor treatment cause *tet2*-mutant HSPC death by inducing apoptosis

Next, we monitored HSPCs as they migrated into the axial vein and traversed to the CHT region in embryos treated from 1 to 3 dpf. At 3 dpf *tet2*−/− HSPC numbers in the CHT region in DMSO-control-treated zebrafish were identical when compared to wild-type zebrafish (**Fig. 3a, top row, and 3b, on the left)**. By contrast, in both selinexor- and eltanexor-treated embryos, the numbers of HSPC in the CHT were significantly decreased in *tet2*−/− compared to wild-type embryos (**Fig. 3a and b).**To determine the cause of lower HSPC numbers at 3 dpf in *tet2*-mutant embryos treated with each drug, we tested the cells for evidence of apoptosis. We performed TUNEL assays coupled with immunohistochemistry (IHC) for GFP in the Tg(*c-myb*:EGFP) reporter line, after treating *tet2*+/+, +/− and −/− embryos with XPO1 inhibitors or DMSO from 1 to 2.5 dpf. We observed a significant increase in the numbers of TUNEL+, GFP+ apoptotic HSPCs in the CHT of *tet2*+/− and *tet2*−/− embryos treated with XPO1 inhibitors, but not in wild-type embryos treated with the same drugs (**Fig. 4a and b**). Thus the lower numbers of *c-myb+* HSPCs at 3 dpf in *tet2*-mutant embryos treated with XPO1 inhibitors observed in Figure 3 were due to the selective induction of apopotosis in *tet2*-mutant HSPCs. We quantified these findings by counting the HSPCs in the CHT region that were double-positive for GFP (*c-myb*) and TUNEL expression using 10 embryos per genotype (**Fig. 4a and b**). We also assessed the possible effects of the drugs on the proliferative activity of the HSPCs by testing the phosphorylation of histone H3 (PH3), which is a marker of cells in mitosis, using the same reporter lines exposed to XPO1 inhibitors or DMSO in the fish water from 1 to 2.5 dpf (**Fig. 1**). These results show that the XPO1 inhibitors do not affect the mitotic index of HSPCs regardless of the genotype, indicating that the drugs did not reduce the fraction of proliferating c-*myb*+ HSPCs in the CHT. Thus, our results indicate that the reduction of *tet2*-mutant HSPC numbers observed after treatment from 1 to 3 dpf by the XPO1 inhibitors is primarily due to increased apoptotic cell death caused by the drugs, without an arrest of cell proliferation.

**Figure 3.**
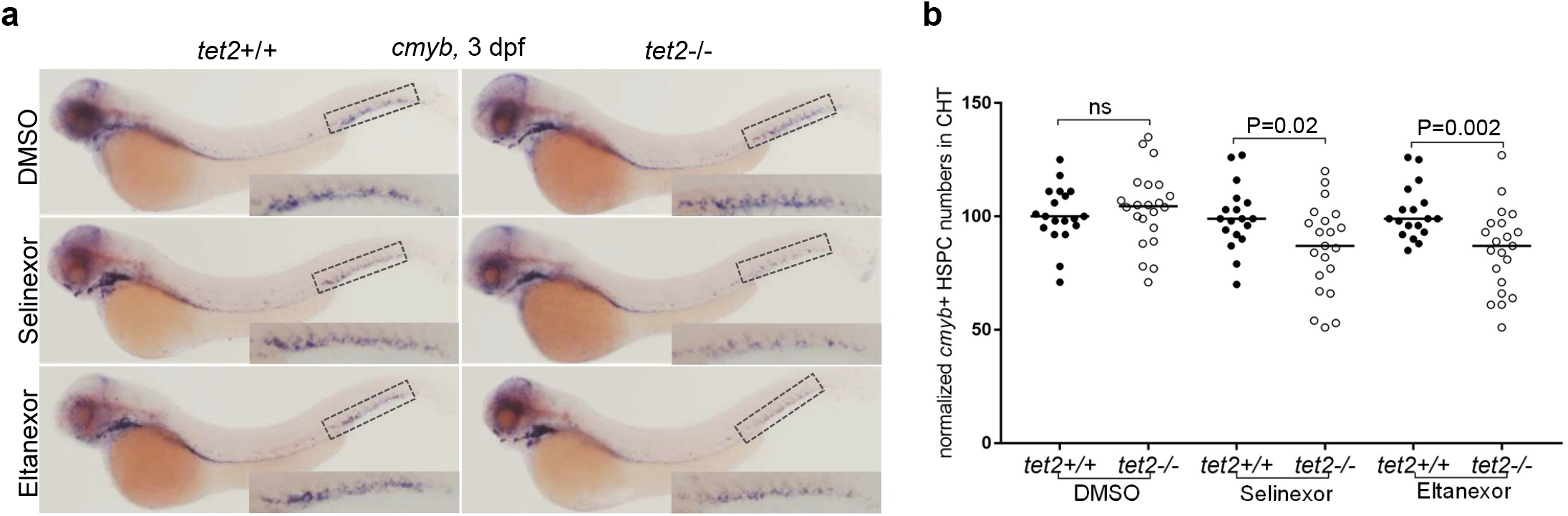
XPO1 inhibitors begin to induce decreased tet2-mutant HSPCs in the CHT region at 3 dpf. **(a)** Embryos with the indicated genotypes were treated with either DMSO, selinexor (100 nM), or eltanexor (250 nM) dissolved in fish water from 1 to 3 dpf. WISH was performed using a riboprobe for *c-myb* at 3 dpf and images were taken. Boxes in each of the images magnify staining in the CHT region. (**b**) Quantification of *c-myb* expression in the CHT region as described in panel a.

**Figure 4.**
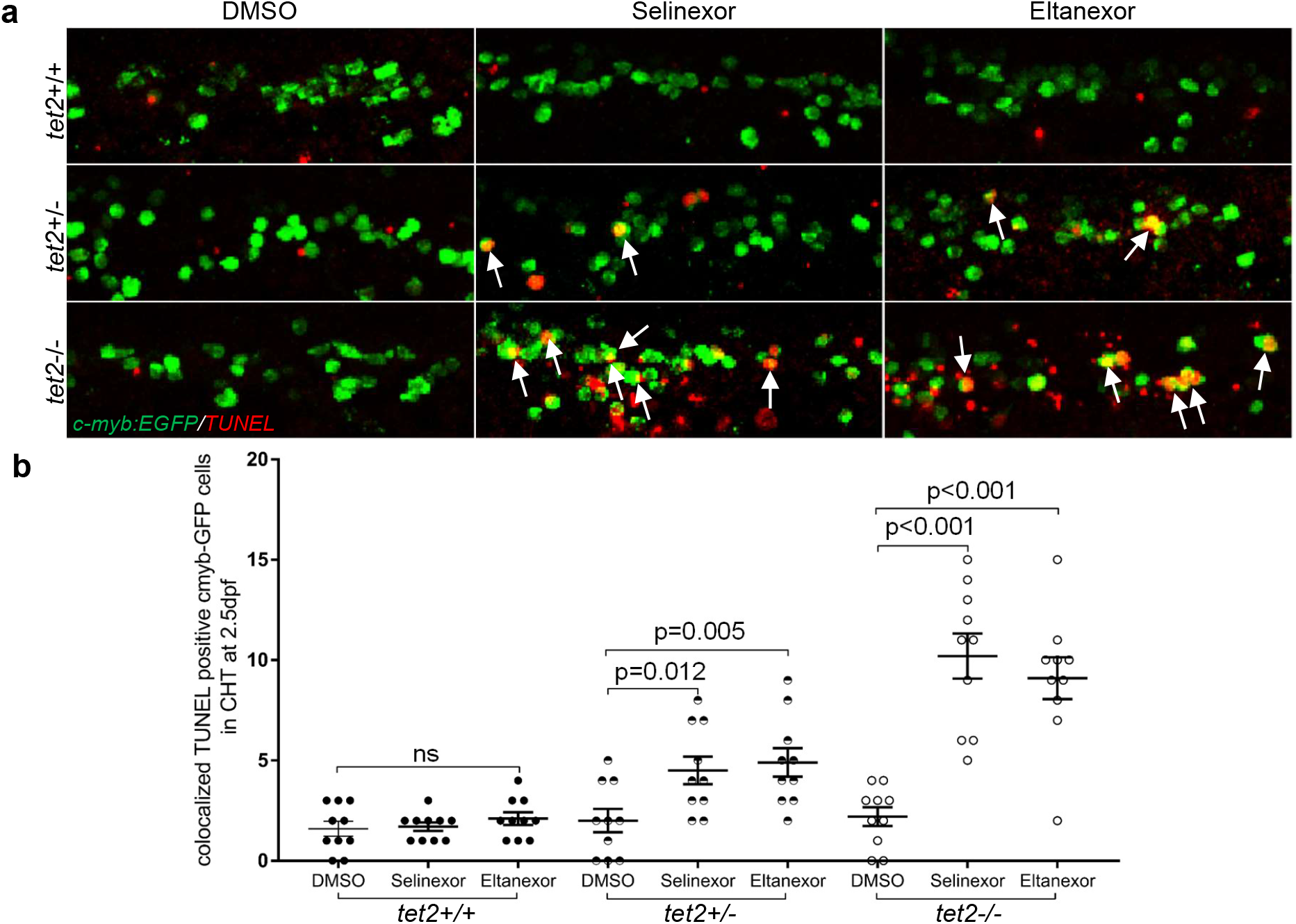
XPO1 inhibitors treatment induces apoptosis in *tet2*-mutant zebrafish HSPCs. (**a**) TUNEL and immunolabeling were performed on 2.5 dpf embryos (treated with either DMSO, 100 nM selinexor, or 250 nM eltanexor) with the indicated genotypes in the *Tg(c-myb:EGFP*)background to identify HSPCs (anti-EGFP antibody; green) and apoptotic cells (TMR-red; red) in the CHT. Fluorescence was visualized using confocal microscopy. White arrows indicate HSPCs undergoing apoptosis. (**b)**Quantification of EGFP+; TUNEL+ cells as described in panel a.

### Selinexor and eltanexor treatment did not induce double-strand DNA breaks in *tet2*-mutant and wild-type HSPCs

In our previous study, TOP1-targeted drugs were synthetic lethal in *tet2*-mutant HSPCs because a deficiency of TDP1 led to reduced levels of repair of single-strand TOP1-DNA adducts, resulting in double-strand DNA damage leading to apoptotic cell death in *tet2*-mutant HSPCs [23]. In evaluating drugs as candidates for treating patients with myeloid malignancies, the induction of double-strand DNA breaks is undesirable, because these patients already harbor mutant HSPCs susceptible to progression to malignancy. Thus, we tested whether XPO1 inhibitors would also cause increased double-strand DNA damage in *tet2*-mutant HSPCs. We performed immunohistochemistry (IHC) using a γH2AX antibody, in 2.5 dpf *tet2+/+* and *tet2*−/− zebrafish embryos generated in the background of the Tg(*c-myb*:EGFP) reporter line to reveal the HSPCs. The *tet2+/+* and *tet2*−/− embryos were incubated from 1-2.5 dpf with selinexor (100 nM) or eltanexor (250 nM), DMSO as a negative control, or topotecan (200 nM) as a positive control. Neither wild-type nor *tet2*-mutant embryos showed detectable γH2AX expression after XPO1 inhibitor treatment, indicating a lack of detectable DNA double-strand breaks in embryos treated with selinexor and eltanexor (**Fig. 5a and b**). As previously observed [23], topotecan as the positive control for this experiment exhibited increased γH2AX expression in *tet2*-mutant HSPCs (**Fig. 5**). This result indicates that the mechanism underlying selective apoptotic cell death following treatment with XPO1 inhibitors in *tet2*-mutant HSPCs does not lead to expression of γH2AX, and thus does not involve increased double-strand DNA breaks.

**Figure 5.**
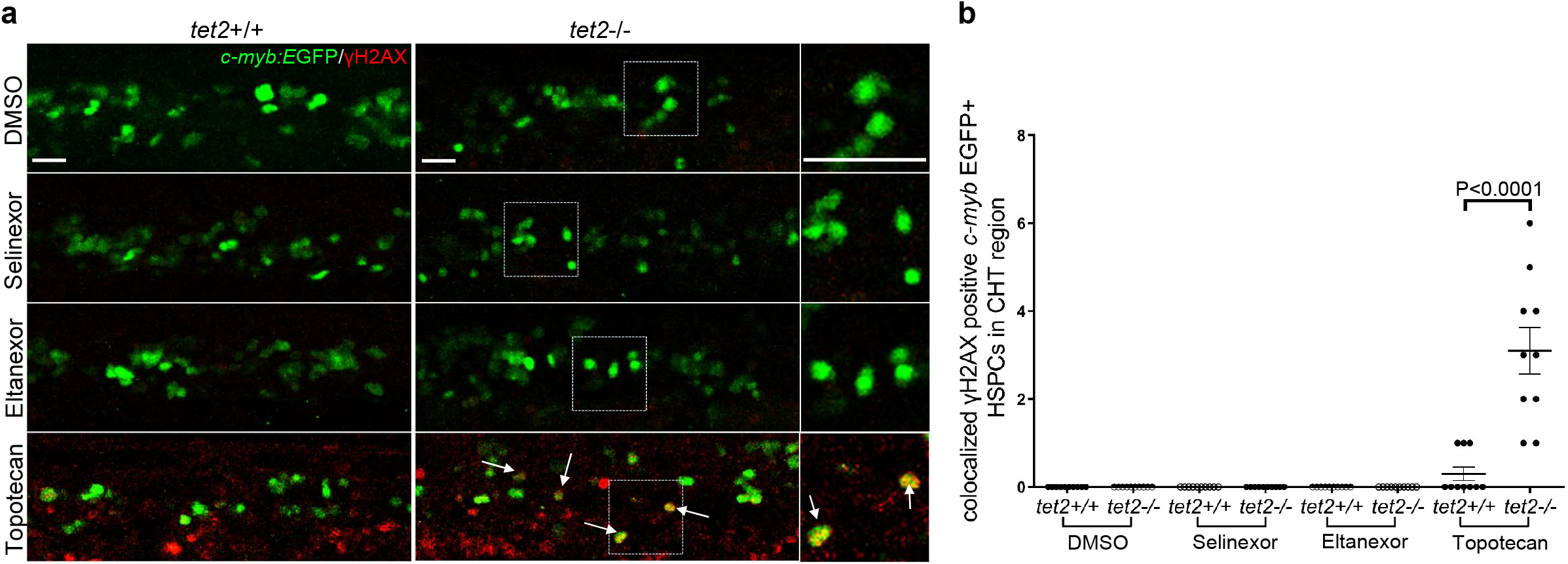
XPO1 inhibitors treatment doesn’t induce DNA double-strand breaks in zebrafish HSPCs. (**a**) Immunolabeling was performed on 2.5 dpf embryos (treated with either DMSO, 100 nM selinexor, 250 nM eltanexor, or 200 nM topotecan) with the indicated genotypes in the *Tg(c-myb:EGFP*) background to identify HSPCs (anti-EGFP antibody; green) and DNA double-strand breaks (anti-γH2AX antibody; red) in the CHT. White arrows indicate groups of HSPCs with DNA double-strand breaks. Magnified views of the regions in the dashed boxes are shown on the right. Scale bar: 25 μm. (**b)** Quantification of EGFP+; γH2AX+ cells as described in panel a.

### Inhibition of XPO1 selectively kills *TET2*-mutant murine HSPCs

To determine whether our zebrafish studies showing a synthetic lethal interaction between *tet2* loss and small molecule inhibition of XPO1 also applies to mammalian HSPC, we first tested the effects of these drugs on the growth of murine *Tet2+/+* and *Tet2*−/− colony-forming HSPCs in methylcellulose medium [27]. For this experiment, the methylcellulose medium supporting colony formation contained either 50 nM selinexor, 50 nM eltanexor, or DMSO control. One thousand LSK progenitors were plated in methylcellulose, and colony formation was quantified after one week (**Fig. 6a**). Addition of XPO1 inhibitors to the methylcellulose dramatically decreased the colony formation of *Tet2*−/− HSPCs, but not of the *Tet2+/+* HSPCs (**Fig. 6b and c**), indicating a selective adverse effect on the growth of *Tet2*-deficient HSPCs compared to wild-type HSPCs.

**Figure 6.**
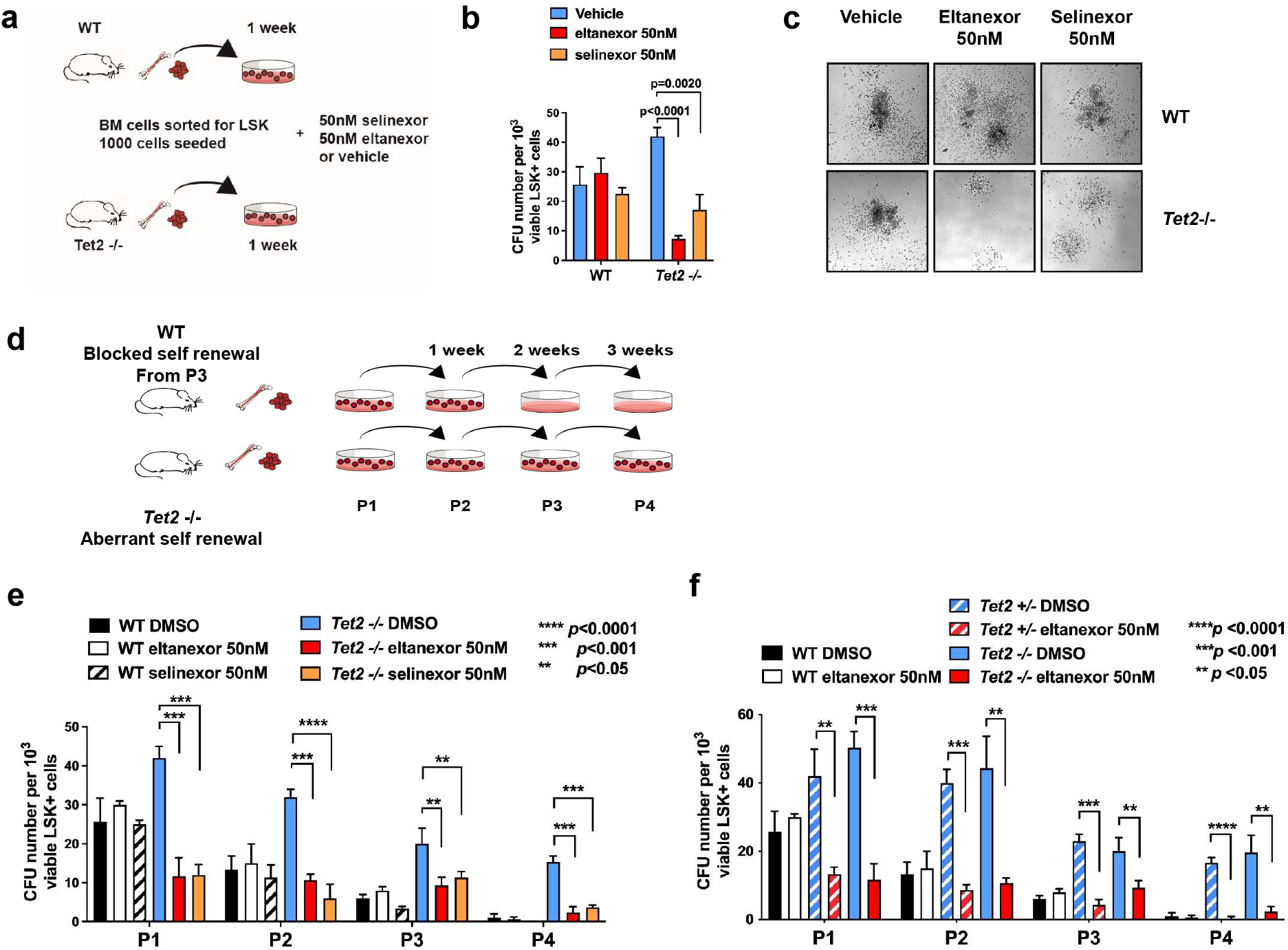
XPO1 inhibitor treatment selectively and dramatically decreases colony formation and the serial replating ability of *Tet2*-mutant murine HSPCs. (**a**) Schematic representation of colony-forming unit assay. BM cells of WT or *Tet2*-mutant mice were sorted for LSK and 1000 LSK+ cells were plated on methylcellulose containing eltanexor, selinexor or vehicle. (**b**) Relative numbers of colonies of murine wild-type or *Tet2*-deficient HSPC cells surviving after treatment with selinexor or eltanexor. The number of colonies was determined after 7 d of culture in methyl-cellulose-containing drug or vehicle. Selinexor and eltanexor reduce *Tet2*-mutant colonies by ~50% or ~75% respectively but have little effect on normal HSPC plating efficiency. (**c**) Representative colony morphology with or without drug treatment. (**d**) Schematic of serial re-plating of murine wild type or *Tet2*-deficient LSK+ HSPC cells in methylcellulose. Cells were treated with selinexor (50nM) or eltanexor (50nM) at each of the four passages (P1-P4). Total number of colony-forming units (CFU) was determined 7d after each passage. (**e-f**) Relative number of colonies of murine wild-type or *Tet2*−/− (e) or Tet2 +/− (f) HSPC cells surviving after treatment with selinexor (e) or eltanexor (e, f) at each of the four passages (P1-P4). Wild-type colonies do not re-plate beyond three passages (P3). *Tet2*-deficient HSPCs exhibit increased replating capacity, which is dramatically decreased by selinexor or eltanexor treatment.

A key feature of *Tet2*-mutant HSPCs in murine models is evidence that loss of *Tet2* imparts an aberrant self-renewal advantage in mutant HSPCs indicated by an increased serial replating capacity in colony-forming assays and a competitive advantage during *in vivo* competitive repopulation studies [10, 27–30]. To determine if XPO1 inhibition reversed aberrant stem cell self-renewal, we performed serial replating assays of *Tet2+/+* and *Tet2*−/− colony-forming bone marrow cells in methylcellulose medium that contained either 50 nM selinexor, 50 nM eltanexor, or DMSO control (**Fig. 6d**). Consistent with previous studies comparing vehicle-treated control *Tet2+/+* and *Tet2*−/− BM LSK+ cells [27], *Tet2+/+* colonies did not replate beyond three passages (P3), while *Tet2*−/− colonies replated for up to four passages (P4), indicating an increased capacity for self-renewal in *Tet2*-mutant HSPC (**Fig. 6e and f**). CFU numbers in each passage were calculated and the results showed that both XPO1 inhibitors induced selective lethality in each passage of *Tet2*-deficient colonies with no effect on wild-type colonies (**Fig. 6e**). We repeated this experiment using eltanexor in *Tet2*+/− BM LSK+ cells, as the heterozygous genotype is often found in humans with MDS and clonal hematopoiesis [14]. Similar to the results we observed in selinexor-treated *Tet2*−/− colonies, eltanexor was selectively lethal to *Tet2*+/− colonies at concentrations of 50nM, in that this drug concentration did not affect *Tet2+/+* colonies (**Fig. 6f**).

### Human AML cell lines with engineered *TET2*-inactivating mutations are sensitive to XPO1 inhibitors

To test whether *TET2*-mutant human AML cells are also hypersensitive to XPO1 inhibitors, we generated loss of function mutations in the *TET2* gene in human MV4-11 cells and K562 cells using CRISPR-Cas9-mediated gene editing. Transduction was performed using lentiviral vectors containing *Cas9* and one of two different CRISPR guide RNAs targeting different sequences within *TET2* exon 3, followed by positive selection with puromycin. Representative clones were isolated from each cell line transduced with the different CRISPR guide RNAs that had biallelic disruption of the *TET2* open reading frame as confirmed by western blotting (clones transduced with the different CRISPR guides are designated as *TET2* mutant #1 and #2, one from each guide RNA, as shown in **Fig. S2**). *TET2* wild-type and *TET2*-mutant MV4-11 and K562 clones derived from each targeting sequence were then cultured with different concentrations of selinexor or eltanexor for 3 days (MV4-11 cells) or 6 days (K562 cells). Cell viability measurements indicated that the *TET2*-mutant clones were more sensitive to both XPO1 inhibitors than the *TET2* wildtype clones (**Fig. 7a, 7b, S3**). We determined that the increased cell growth inhibition induced by XPO1 inhibitors in *TET2*-inactivated compared to wild-type MV4-11 cells was due to measurable apoptosis in the *TET2*-inactivated clones, demonstrated by increased levels of cleaved PARP and cleaved Caspase-3 in *TET2*-inactivated but not wild-type MV4-11 cells by Western blotting (**Fig. 7c)**. We also tested the effect of XPO1 inhibitors on the clonogenic potential of both *TET2*-WT and -mutant MV4-11 cells. In this assay, 1000 cells were seeded in methylcellulose containing DMSO control or 20 nM selinexor or eltanexor. After 7 days in culture, we found that addition of 20nM selinexor or eltanexor to the methylcellulose did not significantly reduce the numbers of colonies for *TET2*-WT MV4-11 cells compared with DMSO treatment. By contrast, selinexor or eltanexor treatment dramatically reduced the number of *TET2*-mutant MV4-11 colonies (**Fig. 7d**).

**Figure 7.**
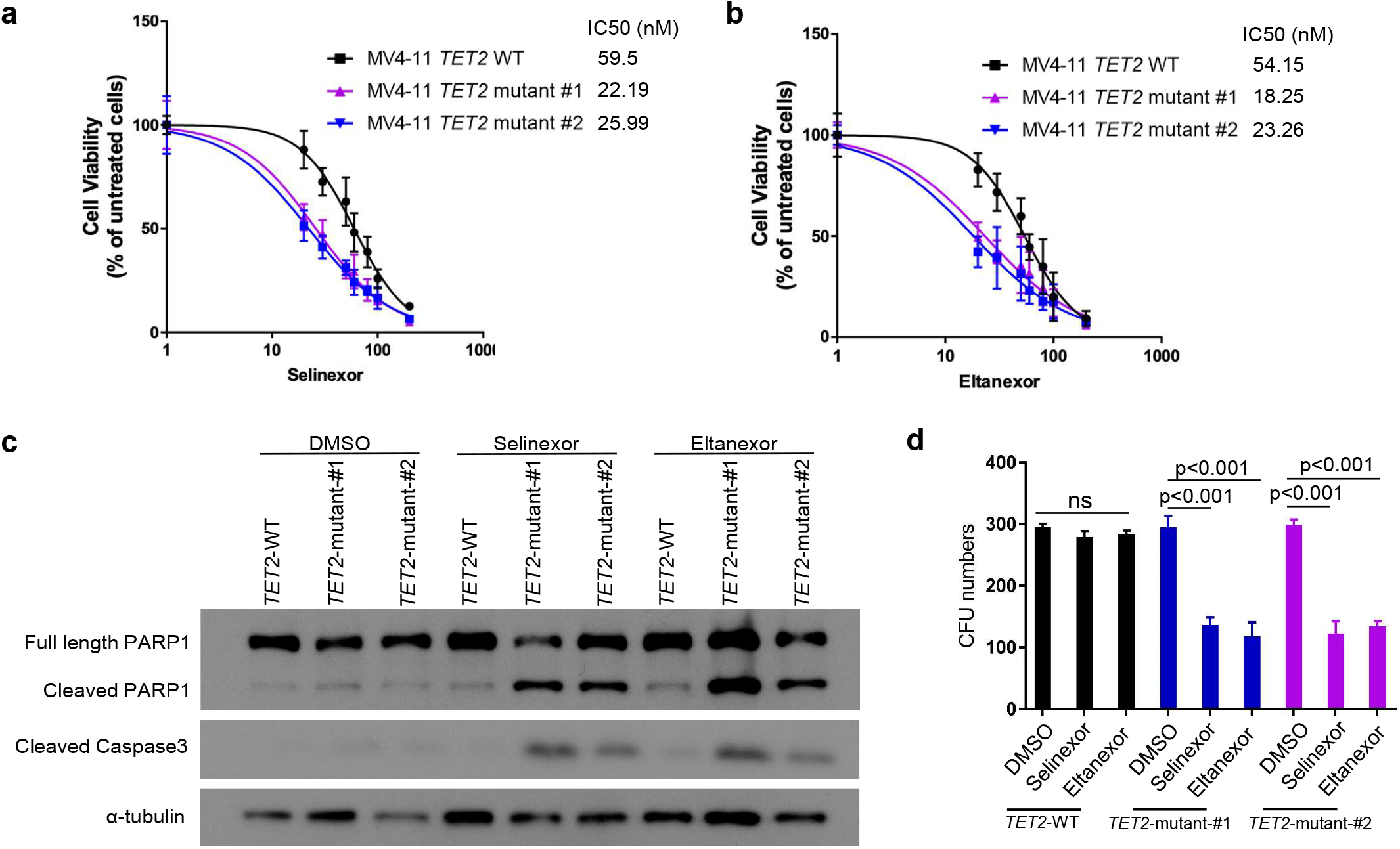
XPO 1 inhibitors selectively kill *TET2*-mutant MV4-11 cells. (**a, b**) *TET2* wild-type MV4-11 leukemia cells and two clones created using different CRISPR guides of CRISPR-cas9-induced, *TET2*-inactivated MV4-11 cells were cultured with different concentrations of selinexor (a) or eltanexor (b) for 72 h. Cell viability was measured with the Cell-Titer Glo assay and the inhibitory concentration at 50% (IC50) is indicated for each group. Data represent the mean SD of three independent experiments. (**c**) *TET2* wild-type and -mutant MV4-11 clones were treated with DMSO or 20 nM selinexor or 20 nM eltanexor for 16 h (for cleaved PARP and cleaved-Caspase3) and analyzed by western blot. This experiment was performed three times with qualitatively similar results. (**d**) *TET2* wild-type and two independent clones of CRISPR-cas9-induced, *TET2*-mutant MV4-11 cells were plated on methylcellulose containing DMSO, selinexor (20nM) or eltanexor (20nM). The total number of colony-forming units was determined after seven days.

## Discussion

Here we report that the XPO1 inhibitors, selinexor and eltanexor, are synthetic lethal with *TET2* inactivation in three different model systems: i) living zebrafish definitive HSPCs treated with the inhibitors *in vivo, ii*) murine bone marrow cell progenitors treated *in vitro* and assayed using colony-forming assays in methylcellulose, and iii) human AML cell lines treated *in vitro*. Taken together, these findings suggest that these XPO1 inhibitors should have a favorable therapeutic index in patients with malignant *TET2*-mutant myeloid leukemia cells compared to the normal bone marrow cells. For example, in our study, XPO1 inhibitors caused increased apoptosis in *tet2*-mutant but not wild-type HSPCs in the zebrafish and *TET2*-inactivated human AML cells, accounting for the enhanced sensitivity of cells lacking *TET2* to this class of drugs (**Fig. 1, 4, and 7**).

### XPO1 mediates the transport of selected proteins and RNAs from the nucleus to the cytoplasm

XPO1, a member of the karyopherin β family, is a major eukaryotic nuclear-cytoplasmic transporter that mediates the transport of a subset of proteins and selected RNA molecules from the nucleus to the cytoplasm [31, 32]. XPO1 regulates the nuclear export of proteins that contain leucine-rich, nuclear-export signals (LR-NES), including protein adaptors that transport RNA molecules [32]. Nuclear export by XPO1 is initiated by regulator of chromosome condensation 1 (RCC1), which is the Ran guanine exchange factor (RanGEF) in the nucleus that converts Ran-GDP into Ran-GTP. Ran-GTP binds to XPO1 in the nucleus, opening XPO1 to bind the NES of cargo proteins, followed by XPO1 transport of cargo molecules through the nuclear pore, followed by Ran-GTP hydrolysis to Ran-GAP with the release of the cargo protein in the cytoplasm [33]. XPO1 cargoes comprise ~1000 eukaryotic proteins, including the tumor suppressor proteins p53, p21, Rb, and FOXO3A, cell cycle regulators, and apoptotic proteins, as well as RNA binding proteins, such as LRPPRC, eIF4E, NXF3, and HuR, along with the RNAs that are transported with them, including RNAs encoding oncoproteins [34, 35].

Expression of XPO1 is upregulated in solid tumors and leukemias, and higher XPO1 levels correlate with a poor prognosis, suggesting an increased dependency of cancer cells on active XPO1-mediated nuclear export [36–38]. As suggested by these high expression levels, nuclear-cytoplasmic transport by XPO1 is required for the survival of several types of solid tumors and hematologic malignancies. Importantly, intermittent drug-mediated inhibition of XPO1 appears to be tolerated by non-neoplastic cells, including normal hematopoietic progenitor cells and proliferating cells of the gastrointestinal tract, thus providing a therapeutic index for XPO1 inhibitors [38, 39].

### Selinexor and Eltanexor are XPO1 inhibitors that are under evaluation as drugs to target hematopoietic malignancies and solid tumors

Phase I studies demonstrated activity of oral selinexor in inducing responses at tolerated doses, including complete remissions in a subset of relapsed refractory multiple myeloma (RRMM), diffuse large B-cell lymphoma (DLBCL) and AML patients. Selinexor was granted accelerated approval by the U.S. Food and Drug Administration (FDA) for the treatment of adult patients with RRMM and relapsed or refractory DLBCL [40–43].

A second-generation XPO1 inhibitor called eltanexor has been introduced, which has a very similar mechanism of action to selinexor but differs in its ability to penetrate the blood-brain barrier [44, 45]. Selinexor is permeable to the blood-brain barrier, which is a benefit when treating tumors of the CNS but leads to CNS-mediated toxicity in the form of anorexia, malaise, and nausea. Eltanexor does not efficiently penetrate the blood-brain barrier, which ameliorates its CNS-mediated side effects compared to selinexor. Thus, eltanexor is currently being evaluated for the treatment of cancers like AML, RRMM and MDS that generally do not involve the brain.

### XPO1 inhibitors do not kill cells by inducing DNA damage

Importantly, we found that XPO1 inhibitors do not cause DNA damage in *tet2-*mutant HSPCs in the zebrafish CHT, based on experiments showing that the XPO1 inhibitor treatment does not cause activation of γH2AX (**Fig. 5**). This is in marked contrast to our earlier results, showing that two other selective drugs for *TET2*-mutant HSPCs, PARP inhibitors and topoisomerase 1-targeted drugs, cause significant DNA damage [23]. Our findings are consistent with the known activity of eltanexor and selinexor in covalently binding to a cysteine in the binding pocket used by XPO1 to engage client proteins for transport from the nucleus to the cytoplasm of the cell. Thus, the lack of DNA breaks in cells treated with eltanexor and selinexor is consistent with their mechanism of action, in marked contrast to PARP inhibitors and topoisomerase 1 targeted drugs, which act by inhibiting the repair of single-strand DNA breaks and allowing them to be converted to double-strand DNA breaks as cell progress into the DNA synthesis phase of the cell cycle. This is an important distinction because bone marrow cells of patients with myeloid malignancies harbor mutations in genes like *TET2* and *DNMT3A* that predispose them to secondary myeloid malignancies induced by treatments that cause double-strand DNA damage.

### Possible mechanisms of increased sensitivity of *Tet2*-mutant HSPCs to XPO1 inhibitors

One hypothesis to explain the increased sensitivity of AML cells and other types of malignant cells is that malignant cells are ‘primed’ to undergo apoptosis [46], so that inhibition of XPO1 by selinexor or eltanexor provides additional apoptotic signals that are sufficient to induce apoptosis in leukemic, but not normal blood cells. However, this mechanism is unlikely to explain the increased sensitivity of Tet2-mutant compared to normal HSPCs, because while Tet2-deficient HSPCs acquire clonal dominance and suppress the growth of normal HSPCs, they are pre-malignant, so that increased apoptotic priming seems unlikely to be linked to increased levels apoptosis. A more likely mechanism in *Tet2*-deficient HSPCs is based on the recently described effect of loss of function of *Tet2* in reducing the expression of key tissue-specific genes. Tet2 preferentially binds to open chromatin in enhancer regions, where it normally demethylates DNA and alters the local chromatin environment to promote the binding and activity of key tissuespecific transcription factors [3]. One hypothesis is XPO1 inhibition in cells with loss of function of Tet2 could lead to even greater reduction in cytoplasmic levels of XPO1 client proteins, because the mRNA expression levels are already reduced without the positive effect of Tet2 on tissuespecific enhancers. Thus in HSPCs with loss of function of Tet2, the levels of cell survival-promoting proteins in the cytoplasm of *Tet2*-mutant HSPCs may fall below critical levels and trigger the expression of BH3-only apoptosis-inducing proteins [46], which mediate cytochrome C release from mitochondria and cause increased levels of apoptosis. We are testing candidate genes expressed at lower levels to see if correction to normal levels in *Tet2-*mutant cells can reverse the aberrant induction of apoptosis that occurs in *Tet2*-mutant HSPCs.

### *TET2* mutations in preleukemic HSPCs and in AML patients in remission

Mutations in *TET2* occur as the “first transforming event” in blood stem cells in individuals who eventually develop MDS and AML and are associated with poor prognosis in AML [6, 8, 12, 47]. It is now recognized that *TET2*-mutant cells persist in AML patients who enter remission, and that the increased self-renewal of these preleukemic cells leads to the acquisition of new *FLT3* and *NPM1* mutations leading to relapse [7, 17]. Thus, there is a need to prevent relapse in AML patients harboring *TET2* mutations after they have achieved remission by targeting these persistent preleukemic *TET2*-mutant cells. ARCH is known to occur in 2-6% of individuals over 50 years of age due to mutations in *TET2* or genes encoding other epigenetic regulatory proteins [48]. These individuals not only have a higher risk of hematopoietic malignancies (preleukemia) but also are at higher risk of cardiovascular disease with an increased risk of heart attacks and strokes, due to the accumulation of mutant macrophages in the vascular wall [14, 15]. Detection of *TET2*-mutant hematopoietic clones in ARCH provides the ideal opportunity to eliminate these preleukemic clones before they acquire additional mutations and progress to AML or cause atherosclerosis with heart attacks and strokes. Our current paper suggests that the XPO1 inhibitor eltanexor can specifically and safely target pre-leukemic, *TET2-*mutant hematopoietic stem and progenitor cells in ARCH to prevent the accumulation of additional mutations with transformation to MDS or AML. There is currently no approved treatment to reduce the levels of *Tet2*-mutant subclones in affected individuals. Further preclinical studies are warranted with the XPO1 inhibitor eltanexor in preclinical *in vivo* models, to clarify the mechanism behind its selective activity in targeting *Tet2*-mutant cells and eltanexor’s potential role in the treatment of patients harboring *TET2* mutations in HSPCs.

## Materials and Methods

### Zebrafish maintenance

All zebrafish lines used in this paper are AB-strain fish and were raised and maintained according to standard procedure. All zebrafish experiments were performed in accord with Dana-Farber Cancer Institute IACUC protocol #02-107. Zebrafish embryos were cultured in “egg water” consisting of 0.03% sea salt and 0.002% methylene blue as a fungicide. The *tet2* mutant zebrafish line was described previously [22].

### Study Selinexor and Eltanexor effect on HSPCs with whole-mount in *situ* hybridization (WISH)

The *tet2*+/−, *tet2*−/− and *tet2*+/+ embryos were incubated with selinexor, eltanexor or DMSO from 1 to 5 days post fertilization (dpf). At 5 dpf, the embryos will be fixed with 4% paraformaldehyde, and effects on HSPCs will be evaluated by whole-mount in situ hybridization (WISH) for the *cmyb*riboprobe. After WISH, the number of HSPCs in the CHT was quantified using ImageJ software, as we have done previously [23].

### Immunofluorescence staining and TUNEL assay

Zebrafish immunofluorescence staining was performed as previously described [23]. Embryos at the indicated stage were fixed with 4% paraformaldehyde at 4°C overnight. For DNA double stand damage study in drug-treated embryos, phospho-gamma-H2A.X (Ser139) antibody (Genetex, GTX127342) and mouse anti-GFP antibody (Thermo Fisher Scientific, Catalog # MA5-15256-D680) were co-incubated at 4°C overnight and visualized with Alexa Fluor 488 goat anti-mouse secondary antibody and Alexa Fluor 546 goat anti-rabbit secondary antibody (Invitrogen). TUNEL apoptosis assay was performed on embryos with the *In-Situ* Cell Death Detection kit (POD: Roche) according to the manufacturer’s recommendations at 37°C for 5 hours. Zebrafish immunofluorescence staining was imaged with a Leica SP5X scanning confocal microscope with a 20x objective. Fluorescence-positive cells were counted in each individual slice, and sum numbers were analyzed with the GraphPad Prism 8 software using the two-tailed Student’s t-test. The optical slice thickness is 3 μm.

### Generation of *TET2*-inactivated AML cells with CRISPR/Cas9

CRISPR-Cas9 technology was used to knockout TET2 in MV4-11 and K562 human leukemia cell lines. PX458 plasmids (pSpCas9-2A-GFP, Addgene, #48138) containing control gRNA (5’-GACCGGAACGATCTCGCGTA-3’) or either of two different gRNAs targeting exon 3 coding sequences of human *TET2* (*TET2*-gRNA #1: 5’-TGGAGAAAGACGTAACTTC-3’ and TET2-gRNA #2: 5’-TCTGCCCTGAGGTATGCGAT-3’) were transfected with Nucleofector (Lonza). After transduction, GFP positive single cells were sorted into 96-well plates by flow cytometry and *TET2* knockout clones were identified with western blotting.

### Protein extraction and western blot analysis

Whole-cell lysates were prepared in RIPA buffer (Cell Signaling Tech) with FOCUS™ ProteaseArrest™ (G-Biosciences) and Phosphatase Inhibitor Cocktail Set II (EMD Millipore). Immunoblotting was performed with each of the specific antibodies to TET2 (CST, #45010), cleaved Caspase-3 (CST, #9664), γH2AX (CST, #9718), α-Tubulin (CST, #5335).

### Cell culture, cell viability analysis

MV4-11 and K562 cells were obtained from the American Type Culture Collection (ATCC). These cells were maintained in RPMI-1640 medium (GIBCO) supplemented with 10% fetal bovine serum (Sigma-Aldrich) and 1% penicillin/streptomycin (Invitrogen). Cells were seeded in a 96-well plate at a density of 5000 cells per well and incubated with dimethylsulfoxide (DMSO) or increasing concentrations of inhibitor. Relative cell growth at day 3 was evaluated by CellTiter-Glo Luminescent Cell Viability Assay (Promega, #G7571). The concentration of inhibitor required for 50% inhibition of cell viability (IC50) was determined with GraphPad Prism 8 (GraphPad Software).

### Mouse colony-forming unit (CFU) assays

In our study, *Tet2*-mutant mice were purchased from Jackson laboratory and represent the knockout line generated by Dr. Anjana Rao [49]. All mouse experiments were carried out in the Dana-Farber Cancer Institute Animal Facility and Lurie Family Imaging Center according to the approved IACUC protocols #13-057. Serial plating assay was described as previously [23]. Murine LSK cells (Lin^-^,Sca-1^+^, c-kit^+^) from 6-8 weeks *Tet2+/+, Tet2 +/−* and *Tet2−/−* mice were purified by flow cytometry and plated in methylcellulose medium (Methocult M3434, Stem Cell Technologies). Total number of colony-forming units (CFU) from *Tet2*+/+, *Tet2 +/−*, and *Tet2* −/− cells in methylcellulose were counted after 7 days. Colonies were isolated and continuously replated for three more passages to compare the serial replating ability between HSPCs from drug and control. Every 7 days, colonies were counted using a Nikon microscope.

### Statistical analysis

Statistical analysis was performed with Prism 5 software (GraphPad). In Figs. 1, 2 and 3, black bars represent the median values from one representative experiment, and the experiment was independently performed at least three times with similar results. In Figs. 4 and 5, bars represent the mean and SEM for 10 embryos from each group from one representative experiment, and experiments were independently performed at least three times with similar results. In Figs. 6 and 7, data are mean ± s.d. of three independent experiments. In Figs. 1,2, 3, 4, 5, 6 and 5, statistical significance was determined using an unpaired Student’s t-test. ns; not significant.

## Authorship contributions

C.-B. J., N.P., Y.L., and A.T.L. designed research; C.-B. J., N.P., performed research; C.-B. J., N.P., M.Z., S.H., Y.L., and A.T.L. analyzed data; and C.-B. J., N.P., M.Z., S.H., Y.L., and A.T.L. wrote the paper.

## Acknowledgments

We thank Cicely Jette and John Gilbert for editorial assistance and critical comments. This work was supported by Edward P. Evans Foundation (A.T.L.), NIH grant R35CA210064 (A.T.L.), the Andrew McDonough B+ Foundation (C.-B.J.), and an International Award of Lady Tata Memorial Trust (C.-B.J.). N.P. was supported by a grant from the Lymphoma Research Foundation.

## Disclosure of Conflicts of Interests

A.T.L. is a founder and shareholder of Light Horse therapeutics, which is pursuing activities unrelated to the current manuscript. Y.L. is an employee of Karyopharm Therapeutics Incorporated and receives compensation and holds equity in the company. All other authors have no competing or financial interests to declare.

---

**Supplementary Figure 1.**
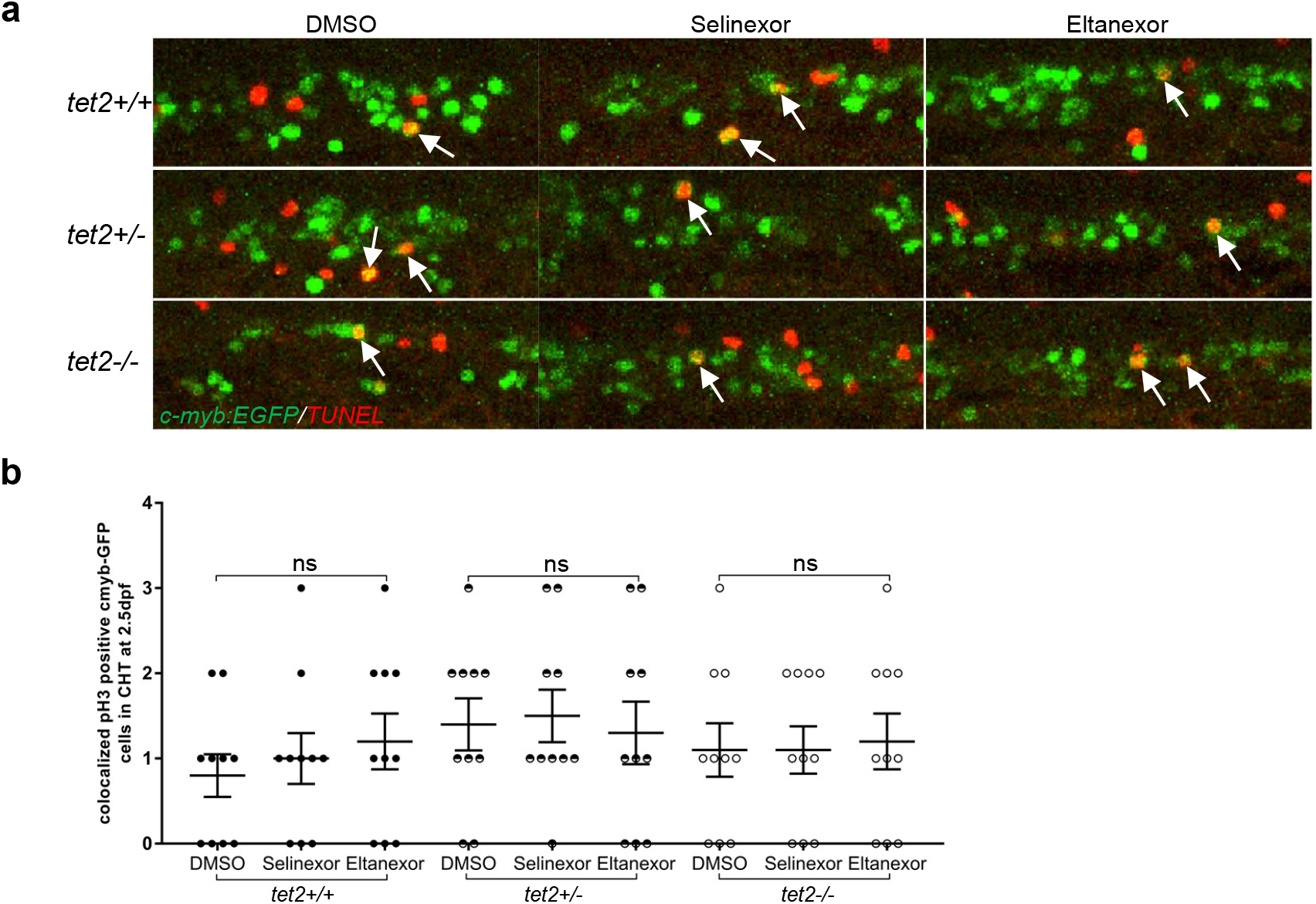
XPO1 inhibitors did not affect the proliferation of tet2 +/+, +/−, and −/− HSPCs. (**a**) TUNEL and immunolabeling were performed on 2.5 dpf embryos (treated with either DMSO, 100 nM selinexor, or 250 nM eltanexor) with the indicated genotypes in the *Tg(c-myb:EGFP*)background to identify HSPCs (anti-EGFP antibody; green) and apoptotic cells (TMR-red; red) in the CHT. Fluorescence was visualized using confocal microscopy. White arrows indicate HSPCs undergoing apoptosis. (**b)**EGFP+; TUNEL+ cell numbers in the CHT region were quantified using ImageJ software. Bars represent the mean and SEM for 10 embryos from each group from one representative experiment. This experiment was independently performed three times with similar results. Statistical significance was determined using an unpaired Student’s t-test. ns; not significant.

**Supplementary Figure 2.**
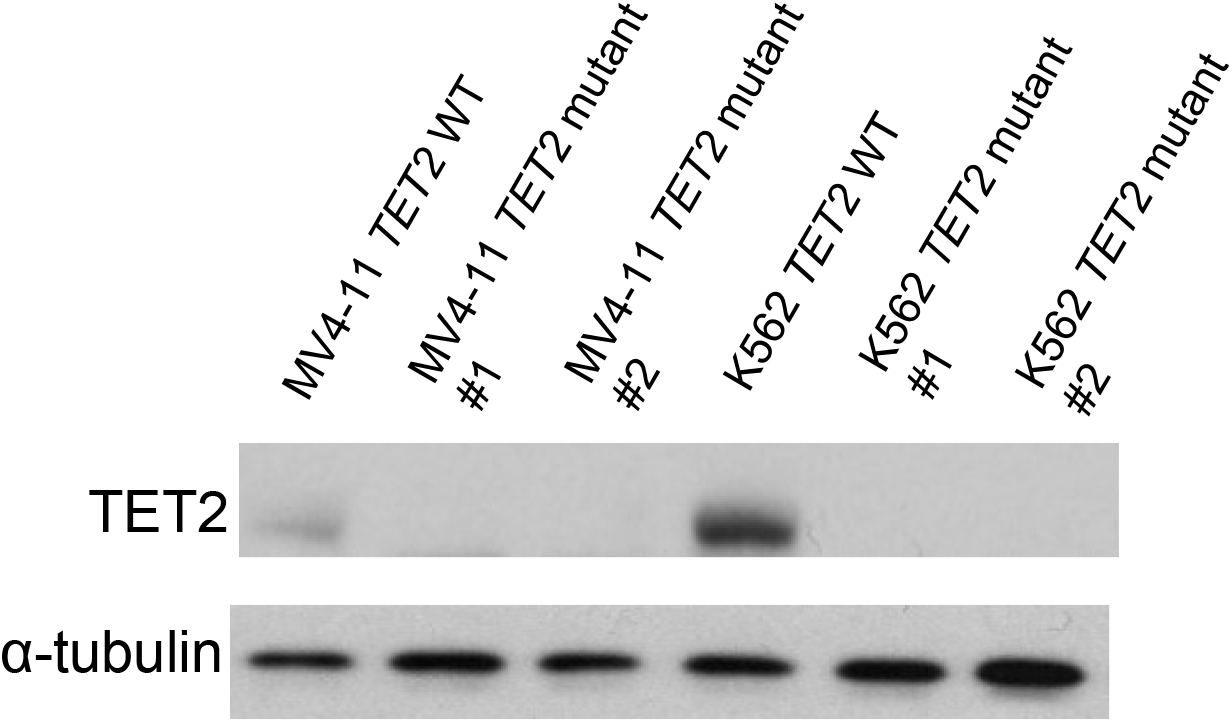
CRISPR-cas9 induced TET2-loss in human AML cells. Two human AML cell lines, MV4-11 and K562, were analyzed for TET2 expression in the *TET2* wild-type parental line and two independent clones of CRISPR-cas9-induced, *TET2*-inactivated clones using western blot. This experiment was performed three times with qualitatively similar results.

**Supplementary Figure 3.**
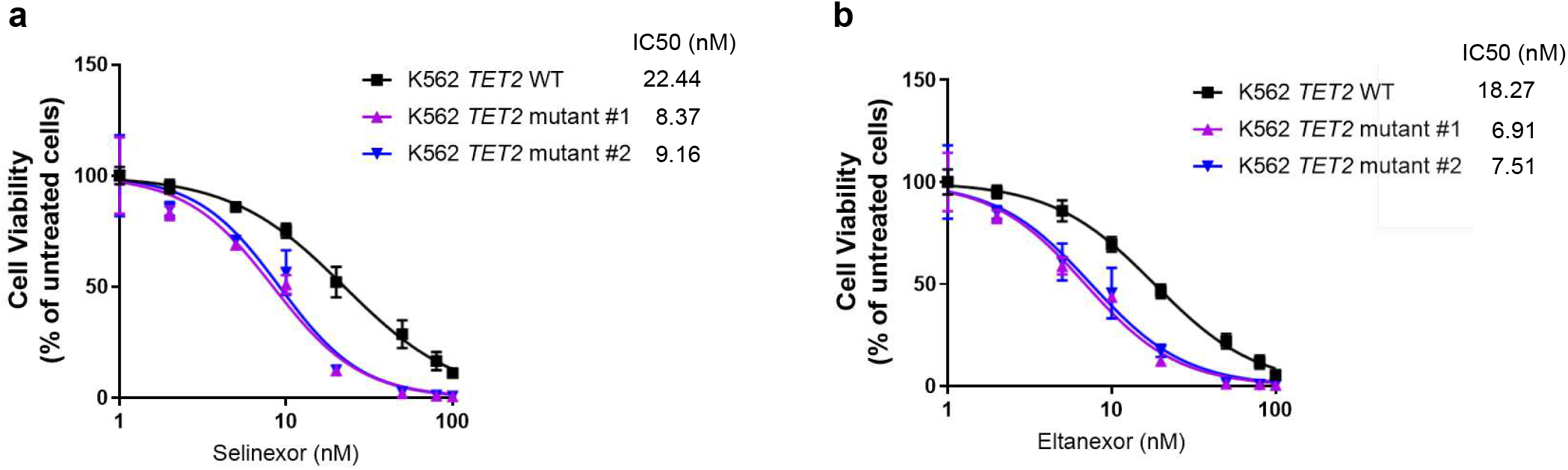
XPO 1 inhibitors selectively kill *TET2*-mutant K562 cells. *TET2* wild-type and two independent clones of CRISPR-cas9-induced, *TET2*-mutant K562 cells were cultured with different concentrations of selinexor (**a**) or eltanexor (**b**) for 6 days. Cell viability was measured with the Cell-Titer Glo assay with the inhibitory concentration at 50% (IC50) indicated for each group. Data represent the mean ±SD of three independent experiments.

